# Febrile temperatures increase *in vitro* antibody affinity for malaria and dengue antigens

**DOI:** 10.1101/345553

**Authors:** Razvan C. Stan, Katia S. Françoso, Rubens P.S. Alves, Luís Carlos S. Ferreira, Irene S. Soares, Maristela M. de Camargo

## Abstract

Fever is a regulated elevation in the body setpoint temperature and may arise as a result of infectious and noninfectious causes. While beneficial in modulating immune responses to infection, the potential of febrile temperatures in regulating antigen binding affinity to antibodies has not been explored. We have investigated this process under *in vitro* conditions using selected malaria or dengue antigens and specific monoclonal antibodies, and observed a marked increase in the affinity of these antibody-antigen complexes at 40°C, compared to physiological (37°C) or pathophysiological temperatures (42°C). Induced thermal equilibration of the protein partners at these temperatures, prior to measurements, further increased their binding affinity. These results may indicate an unexpected beneficial and adaptive role for fever *in vivo*, and highlight the positive role of thermal priming in enhancing protein-protein affinity for samples of scarce availability.

---

Maintaining a constant temperature in mammals is a tightly regulated process, including in cases where infections occur and body temperature increases. Fever, an initial, nonspecific, acute-phase response during infections, is a key factor responsible for improving survival and shorten disease duration (1). Cellular events occurring during physiological fever or hyperpyrexia have been the focus of intensive clinical and *in vitro* studies (2). Previous research has revealed that fever-inducing pathogen load is reduced due to enhanced host defence while its proliferation at febrile temperatures is not significantly affected. (3). Physiological and reversible increases in core body temperatures do not normally exceed the 40°C threshold (4), with survival beginning to decrease when fever exceeds 39.5°C, suggesting that there is an upper limit to the optimal fever range (5).

While elevated temperatures profoundly alter membrane fluidity, cell signalling and gene expression patterns, the role of febrile temperatures in directly affecting antibody affinity for antigens from pathogens that induce fever has not been explored. We have focused here on the *in vitro* changes of binding affinities for two antibody-antigen immune complexes characteristic of two widespread, fever-inducing infectious diseases (6, 7). To this end, we made use of antigens from a viral agent, *i.e.* non-structural protein 1 (NS1) from dengue virus serotype 2 DENV-2 (8), and from a protozoan pathogen, namely the 19-kDa carboxyterminal region of merozoite surface protein 1 (MSP1_19_) from *Plasmodium vivax* (9), and their corresponding monoclonal IgG antibodies (10, 11).

Antibodies progressively mature their affinity and specificity for various target antigens by changing the amino acid residue composition of their complementarity-determining regions (12). As with other proteins, high affinity can be achieved by fast association rates coupled to slow off-rates in a process directly dependent, among other factors, on temperature. The thermal optimum of antibody-antigen complex depends on the chemical nature of the epitope and paratope, and on the type of bonds formed at different temperatures (13). The effect of fever temperature on equilibrium constants of immune complexes formed by erythrocyte antibodies has previously uncovered very modest decreases from 2.2 10^7^ M^−1^ at 15□-19°C to 1.8 10^7^ M^−1^ at 37□-40°C (14).

## Results

### Isothermal Titration Calorimetry (ITC) and ELISA measurements

We carried out with ITC a systematic investigation into the formation of dengue and malaria immune complexes at 37°C, 40°C, and 42°C. Prior to measurements, we have equilibrated *in vitro* at indicated temperatures all the proteins for 1 hour (Supporting Information). This procedure was carried out in order to subject the partner proteins to temperatures more relevant to those caused by fever-inducing pathogens and for periods that *in vivo* are not lethal (e.g. less than 6 hours at 41□-42□) for *P. vivax* infections) (15). Representative ITC thermograms are shown in Figure 1.

**Fig. 1.**
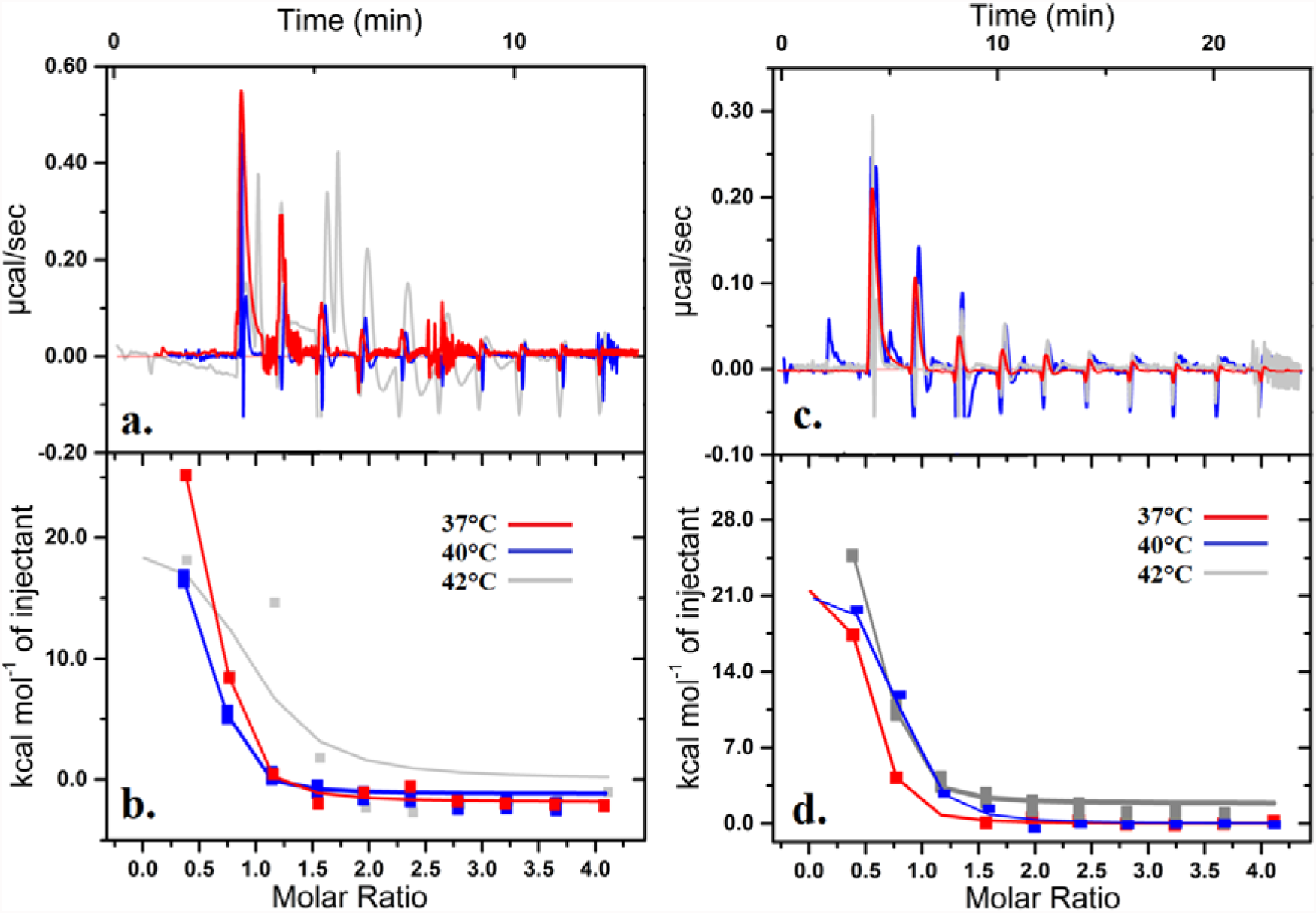
ITC measurements of malaria MSP1_19_ titrated into K_2_3 IgG antibody (a-b) and DENV-2 NS1 titrated into 4H1BC IgG1 antibody (c-d) at indicated temperatures. Results fitted to a single-binding site model with 1:1 stoichiometry.

The changes in enthalpy with increases in temperature are nonlinear, characterized by lowest values at 40°C for both systems, followed by a more negative change at 42°C and a further decrease at 37°C (DENV-2), while the opposite is present for the malaria immune complexes (Figure 1). A common feature that we measured for both systems was an increase in binding affinity at 40°C compared to other temperatures, as measured in solution by ITC, or in solid-phase assay with ELISA, as presented in Figure 2.

**Fig. 2.**
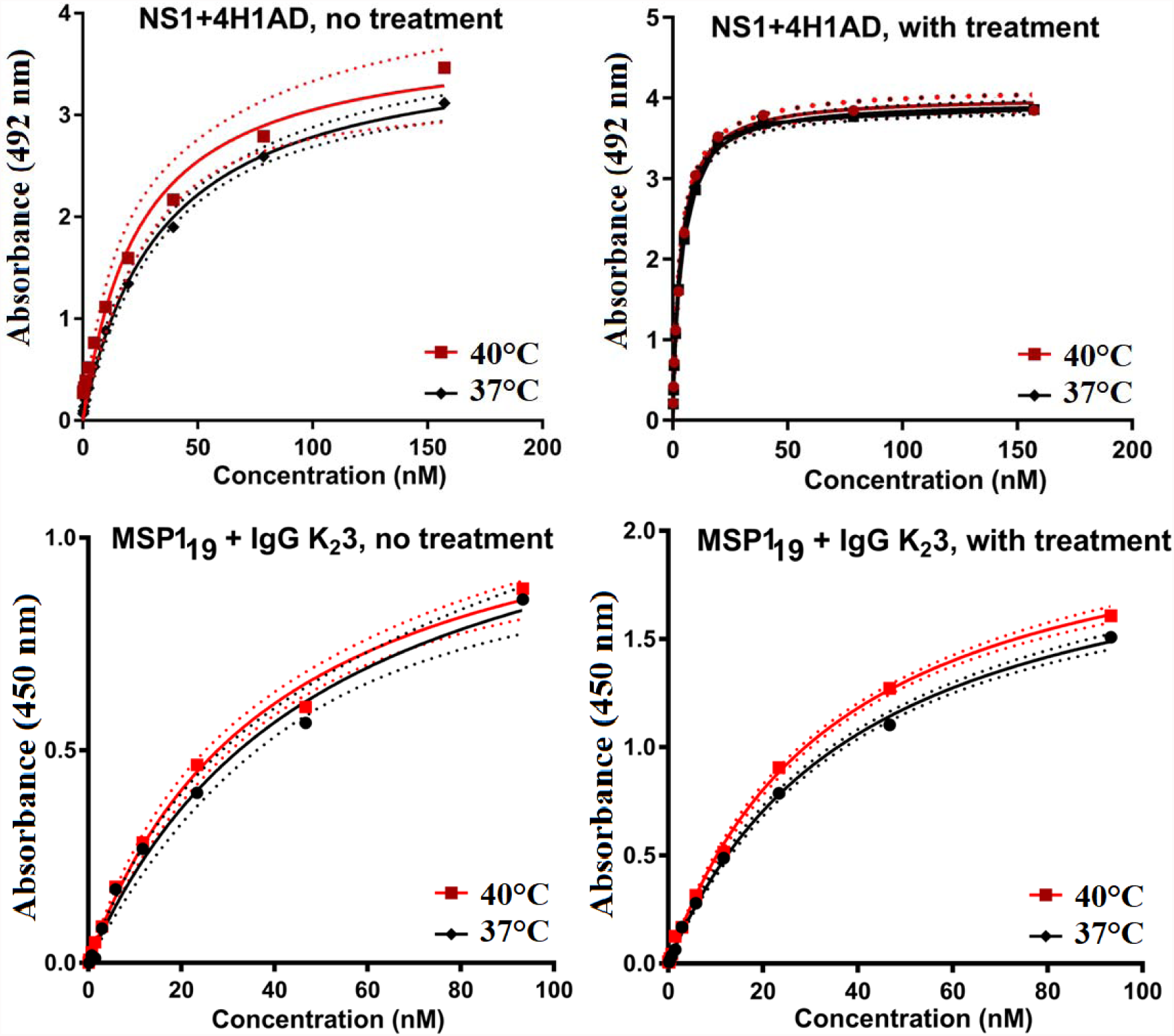
Upper panel: ELISA measurements of dengue DENV-2 NS1 antigen with IgG1 4H1BC, without (a) or with (b) a thermal pre-equilibration step. Lower panel: ELISA measurements of malaria MSP1_19_ antigen with IgG K_2_3, without (c) or with (d) a thermal treatment procedure at indicated temperatures.

For both immune complexes, ELISA measurements showed a ~1.15-1.3 increase in affinity from 37°C to 40°C, in the presence or absence of thermal priming. However, thermal pre-equilibration led to significant improvements in KD by a factor of ~9.5 (at 37°C) and ~8 (at 40°C) for the dengue immune complex, and to a decrease in KD by a 1.3 factor (at 37°C) and a 1.2 factor at 40°C, respectively for the malaria complex. An overview of ITC and ELISA results is shown in Table 1.

**Table 1.**
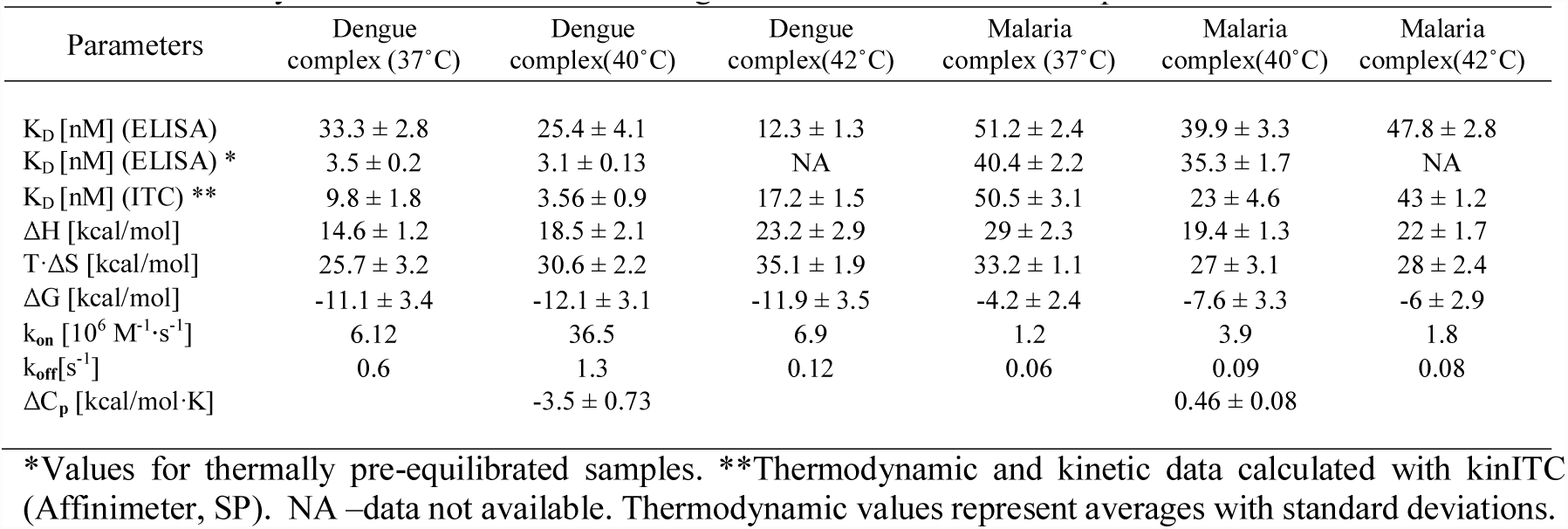
Thermodynamic and kinetic data of dengue and malaria immune complexes

| Parameters | Dengue complex (37°C) | Dengue complex(40°C) | Dengue complex(42°C) | Malaria complex (37°C) | Malaria complex(40°C) | Malaria complex(42°C) |
| --- | --- | --- | --- | --- | --- | --- |
| K<sub>D</sub> [nM] (ELISA) | 33.3 ± 2.8 | 25.4 ± 4.1 | 12.3 ± 1.3 | 51.2 ± 2.4 | 39.9 ± 3.3 | 47.8 ± 2.8 |
| K<sub>D</sub> [nM] (ELISA) * | 3.5 ± 0.2 | 3.1 ± 0.13 | NA | 40.4 ± 2.2 | 35.3 ± 1.7 | NA |
| K<sub>D</sub> [nM] (ITC) ** | 9.8 ± 1.8 | 3.56 ± 0.9 | 17.2 ± 1.5 | 50.5 ± 3.1 | 23 ± 4.6 | 43 ± 1.2 |
| ΔH [kcal/mol] | 14.6 ± 1.2 | 18.5 ± 2.1 | 23.2 ± 2.9 | 29 ± 2.3 | 19.4 ± 1.3 | 22 ± 1.7 |
| T·ΔS [kcal/mol] | 25.7 ± 3.2 | 30.6 ± 2.2 | 35.1 ± 1.9 | 33.2 ± 1.1 | 27 ± 3.1 | 28 ± 2.4 |
| ΔG [kcal/mol] | -11.1 ± 3.4 | -12.1 ± 3.1 | -11.9 ± 3.5 | -4.2 ± 2.4 | -7.6 ± 3.3 | -6 ± 2.9 |
| k<sub>on</sub> [10<sup>6</sup> M<sup>-1</sup>·s<sup>-1</sup>] | 6.12 | 36.5 | 6.9 | 1.2 | 3.9 | 1.8 |
| k<sub>off</sub> [s<sup>-1</sup>] | 0.6 | 1.3 | 0.12 | 0.06 | 0.09 | 0.08 |
| ΔC<sub>p</sub> [kcal/mol·K] |  | -3.5 ± 0.73 |  |  | 0.46 ± 0.08 |  |

\*Values for thermally pre-equilibrated samples. \*\*Thermodynamic and kinetic data calculated with kinITC (Affinimeter, SP). NA –data not available. Thermodynamic values represent averages with standard deviations.

## Discussion

ITC measurements for the dengue system at 40°C revealed a K_D_ lower by a factor of ~2.8 compared to values at 37°C and correspondingly by a factor of ~5, compared to measurements at 42°C. The main contribution to binding affinity is entropic, with the highest value measured at 42°C, being compensated by large positive enthalpy variations. The intrinsic free energy of binding was largely independent of temperature variations, reflecting the compensation of the enthalpic and entropic terms. Kinetic parameters were largely responsible for the differences in affinities observed across temperatures, with much faster k_on_ rates measured at 40°C, especially when compared to values at 42°C (~11 fold increase). Epitope-mapping studies have identified the sequence ^193^AVHADMGYWIESALNDT^209^ as the conformational epitope buried within a β-sheet structure of the dengue NS1 protein (16). The presence therein of net three negative charges conferred by the aspartic and glutamic acid residues may result in more stable immunocomplexes at higher temperature, as previously measured in other systems (17). The negative grand average of hydropathy indicates a hydrophilic interface for this immune complex, whose hydration on temperature increase lies (18) at the origin of the observed negative heat capacity. Importantly, the effect of temperature changes in modulating monoclonal antibodies binding to DENV-2 antigens was also shown to be important for its structural E proteins (19). Unique quaternary conformational epitopes were exposed when virions were incubated at vertebrate host physiological temperature (37°C), versus the mosquito thermal optimum of 28-30°C (19). ITC measurements for the malaria complex showed a K_D_ approximately ~2.1 and ~1.8 folds lower at 40°C when compared to values at 37°C and at 42°C, respectively. The peak in affinity at 40°C is, unlike the DENV-2 system, due to both high k_on_ rates and to more favorable enthalpic contribution to binding, compared to the measurements at 37°C and at 42°C. While the structure of the antigen-binding site for this malaria immune complex is unknown, our measurements indicate that larger unfavorable entropy values obtained at 37°C and 42°C, compared to results at 40°C. Such effect, coupled with the overall positive heat capacity change of the reaction, are suggestive of predominantly hydrophobic solvation dominated by both unfavorable enthalpy and a consequence of favorable entropy (20).

We have not detected with Circular Dichroism experiments significant secondary structure modifications in any protein partner or complex, across the temperature range here used (Figure S1). The differences in affinities that we measured between the two techniques have previously been reported for a malaria system involving MSP1 (21). These discrepancies may be accounted for by the presence of the adsorbed phase in ELISA measurements that could hinder epitope binding, produce steric or attractive interactions between the mAb molecules (22) themselves or induce blocking of binding sites by multivalent binding and rebinding (23). These restrictions may be reduced for thermally equilibrated ELISA samples, resulting in higher affinities compared to ITC results or non-equilibrated ELISA data (Table 1).

In conclusion, our data indicate the potential for reversible, physiological fever temperatures in augmenting antibody affinity *in vitro* for tertiary and quaternary epitopes. This new role may constitute an important adaptive mechanism for antibody-mediated detection and protection against pathogens. Further validation by *in vivo* studies and extension to a larger set of antigens involved in fever episodes, including from bacterial pathogens, will extend the reach of our conclusions. In addition, our results may add to the growing interest in relating hyperthermia to the efficiency of cancer immunotherapy (24). Finally, active thermal equilibration of protein partners prior to performing ELISA or other relevant *in vitro* assays is proposed to improve on binding affinities and assist in cases where limited amounts of samples are available.

## Methods

### Enzyme-linked immunosorbent assays (ELISA)

#### ELISA measurements with dengue DENV-2 antigen and antibody

ELISA with solid-phase bound NS1 protein were carried out as previously described (8). Briefly, polystyrene Maxisorp microplatesNunc were coated overnight at 37°C, 40°C or 42°C with a purified recombinant NS1 expressed in *Escherichia coli* (400 ng/well) in triplicates. The plates were washed 3 times with phosphate-buffered saline (PBS) containing 0.05% Tween-20 (PBST) and blocked with 1xPBST containing 3% skim milk and 0.1% of BSA for 2 hours at 37°C or 40°C. After a new wash cycle, the anti-NS1 mAb 4F6 (11) was diluted (log2) starting at 157.3 nM, added to wells and incubated at 37°C or 40°C for 2 hours. After a new wash cycle, the anti-mouse IgG antibody conjugated to peroxidase (Sigma, USA) was added to wells and incubated again for 2 hours at 20 ± 2°C. After a final washing cycle, plates were developed with sodium citrate buffer (pH 5.8) containing ortho-phenylenediamine dihydrochloride (OPD) (Sigma, USA) and H_2_O_2_ and the reaction was stopped after 15 min with the addition of 50 µl of H_2_SO_4_ at 2 M. The optical density reading was performed at 492 nm plate reader (Labsystems Multiscan, Thermo-Scientific, USA).

#### ELISA measurements with malaria PvMSP1_19_ antigen and antibody

Recombinant protein PvMSP1_19_ was kept at 37°C, 40°C or 42°C for 1 hour prior to ELISA assays. The recombinant protein was employed as solid phase-bound antigen (200 ng/well) and a volume of 50 µl of each solution was added to each well of 96-well plates (BD Costar 3590). After overnight incubation at each indicated temperature, the plates were washed with PBST and blocked with 5% milk-2.5% BSA for 2 hours, at each specified temperature. The plates were washed with PBST and the monoclonal antibody K_2_3 (10) was tested at serial dilutions (2x) initiating at 93.32 nM in a final volume of 50 µl of sample added to each well in triplicate, followed by incubation for 1 hour at each temperature. After washes with PBST, 50 µl of a solution containing anti-mouse IgG (KPL) conjugated to peroxidase diluted 1:3.000 was added to each well and incubated at 20±2°C for 1 hour. The enzymatic reaction was developed using 3,3′, 5,5′tetramethylbenzidine (TMB) (Bio-Rad)) for 15 minutes and stopped using H_2_SO_4_ 1N. The optical density values were determined at 450nm.

All ELISA data was analyzed using Prism 7 (GraphPad, USA). Background-subtracted data represent averages of two to three independent readings.

#### Circular dichroism (CD)

CD measurements were performed in a JASCO-810 spectrometer (Jasco, Japan) coupled to a Peltier temperature controller (Model JWJTC-484). Immunocomplexes (0.064 mg/ml MSP1_19_ with 0.256 mg/ml IgG K_2_3, and 0.128 mg/ml NS1 with 0.64 mg/ml IgG 4H1BC, respectively) were reconstituted at 20 ± 2°C. Three hundred µl of each complex were immediately thereafter placed in the 1-mm CD cell for 1 hour at specified temperatures, prior to data acquisition. All measurements were performed in 10 mM PBS, pH 7.4, containing 1 mM 2-mercaptoethanol. Data were averaged from three scans at 100 nm/min, data pitch 0.1 nm, bandwidth 1 nm. Buffer baselines were subtracted from respective sample spectra.

#### Isothermal titration calorimetry (ITC)

Protein concentrations were determined spectrophotometrically by measuring the absorbance at 280 nm (NanoDrop 2000, Thermo-Scientific, USA). Molar absorption coefficients for all proteins were calculated with ProtParam (SIB, Switzerland). ITC measurements were performed on a MicroCaliTC 200 calorimeter (GE Healthcare, USA). The heat signals due toprotein-protein binding were obtained as the difference betweenthe heat of reaction and the corresponding heat of dilution, using Microcal Origin v7.0 (OriginPro, USA). Thermodynamic and kinetic values based on ITC data were obtained using KinITC (25) (Affinimeter, Spain). All titrations comprised of antigen in syringe at 7 µM (NS1) or 1 µM (MSP1_19_) and 10µM (anti-NS1 IgG) or 1 µM (anti-MSP1_19_ IgG) in the cell, respectively. All measurements were performed in 100 mM PBS, pH 7.4, with 1 mM of 2-mercaptoethanol. IgGs were kept in iTC200 cell at each indicated temperature for 1 hour prior to measurements. Antigens were separately heated for the same period in 200 µL Eppendorf microcentrifuge tubes in a water bath (Thermo-Scientific, USA) before start of titrations.

## Supporting Information Available

Circular Dichroism data are included in the supporting information.

## Acknowledgments

We thank Dr. Iolanda M. Cuccovia and Dr. Michael McLeish for critical reading of the manuscript. We appreciate the technical support from Darshak Bhatt and Eneas de Carvalho. This study was funded by the Brazilian National Council for Scientific and Technological Development (400662/2014-0 for R.C.S. and 309041/2012-0 for M.M.D.C.) and São Paulo Research Foundation (2014/17595-0 for L.C.S.F.).

